# Invasion of spontaneous germinal centers by naive B cells is rapid and persistent

**DOI:** 10.1101/2023.05.30.542805

**Authors:** T. van den Broek, K. Oleinika, S. Rahmayanti, C. Castrillon, C.E. van der Poel, M.C. Carroll

## Abstract

In autoreactive germinal centers (GC) initiated by a single rogue B cell clone, wild-type B cells expand and give rise to clones that target other autoantigens, known as epitope spreading. The chronic, progressive nature of epitope spreading calls for early interventions, but the kinetics and molecular requirements for wild-type B cell invasion and participation in GC remain largely unknown. With parabiosis and adoptive transfer approaches in a murine model of systemic lupus erythematosus, we demonstrate that wild-type B cells join existing GCs rapidly, clonally expand, persist, and contribute to autoantibody production and diversification. The invasion of autoreactive GCs required TLR7, B cell receptor specificity, antigen presentation, and type I interferon signaling. The adoptive transfer model provides a novel tool for identifying early events in the breaking of B cell tolerance in autoimmunity.

**One Sentence Summary:** The autoreactive germinal center is an open structure that is susceptible to rapid and persistent naive B cell invasion, with clonal expansion and auto-antibody induction and diversification.

## INTRODUCTION

Systemic lupus erythematosus (SLE) is a prototypical autoimmune disease characterized by a diverse and evolving repertoire of autoantibodies, which are often present prior to the clinical manifestation of disease and drive multi-organ involvement (*1, 2*). In the healthy individual, the mature B cell repertoire contains autoreactive B cell clones, but several tolerance checkpoints prevent them from being activated and producing auto-antibodies (*3, 4*). However, after the initial breakdown of self-tolerance to a single nuclear self-antigen, such as small nuclear ribonucleoproteins (RNPs), there is a diversification of antibody self-specificity, called epitope spreading, that leads to overt disease (*2, 5*).

Germinal centers (GC) are the primary sites for generating high-affinity antibodies and subsequent epitope spreading (*6*). As such, early interventions in the GC response could lead to better therapies. We have previously shown that GCs consist mainly of WT B cells that undergo clonal selection and maturation, resulting in epitope spreading (*7*). What is more, WT B cells eventually become independent of the autoreactive B cells, being able to sustain the GC response in their absence. For that study, we used a mixed bone marrow chimera model, where wild-type (WT) B cells co-exist with transgenic B cells specific for ribonuclear complexes (derived from 564Igi mouse bone marrow) (*7, 8*). However, this model lacks a way to synchronize B cell entry into the GC, so the chronicity of WT B cell engagement precludes the specific evaluation of early mature B cell activation, GC entry, and autoantibody production. Furthermore, apart from TLR7 (*7, 8*), the molecular requirements for WT B cell participation in autoreactive GCs are unknown.

Here, by synchronizing WT B cell entry, via parabiosis and adoptive transfer experiments, we pinpoint when and how naïve WT B cells gain initial access to the autoreactive germinal center. These data indicate that polyclonal naïve B cells can rapidly enter, clonally expand, and persist within spontaneous GC. Notably, via adoptive transfer of WT B cells, we identified crucial factors involved in their invasion of spontaneous GC and hereby provide a tool to further study the requirements for GC entry.

## RESULTS

### WT B cells enter and dominate mature autoreactive GCs

We used a parabiosis model to characterize the overall dynamics of WT B cells with synchronized access to spontaneous mature GCs. Specifically, WT mice were surgically bound to 564Igi or WT mice (control parabiosis) (Fig.1A), and the B and T cell exchange between mice was assessed by periodic blood draws and tracked using allelic variants of the pan-leucocyte marker CD45 (Fig.1B, C). In the 564Igi:WT parabiont, B and T cell exchange in the peripheral blood reached 50% at 2 weeks following parabiosis (Fig.1D, Suppl.Fig.1A). Peripheral blood T cell frequencies equilibrated thereafter at comparable exchange rates between the 564Igi:WT and control WT:WT parabiosis (Suppl.Fig.1A,B), indicating a successful surgery. However, WT B cells began to dominate the circulation of the 564Igi:WT parabiont by two weeks and increased over time (Fig.1D, gray line), thereby reducing the proportion of circulating 564Igi B cells (Fig.1E). We looked beyond the circulation and found that, at 8-9 weeks post-surgery, WT B cells were significantly enriched within the splenic GC compartment (phenotypically GL7+CD38-) of the 564Igi partner; in contrast, this pattern was not observed for 564Igi B cells in the WT parabiont (Fig.1F).

**Figure 1.**
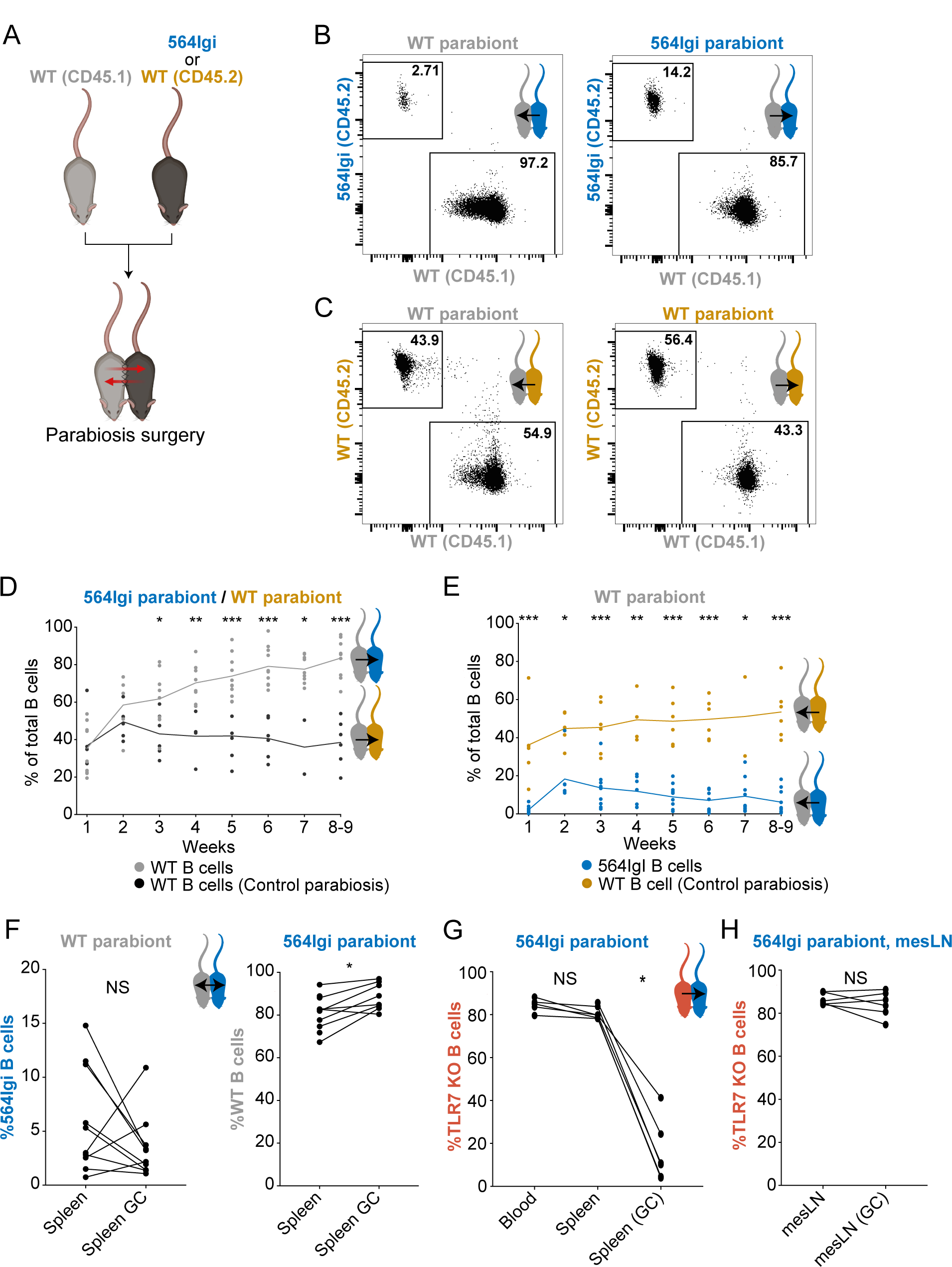
TLR7-dependent wild-type B cell dominance in spontaneous GCs in 564Igi:WT parabionts. A. Schematic depiction of the parabiosis model (WT = wild-type). B. Representative FACS dot plot of the proportion of WT or 564Igi B cells in the circulation of the WT and 564Igi parabionts, at 6 weeks parabiosis. C. Representative FACS dot plot of the proportion of WT (CD45.2) or WT (CD45.1) cells in the circulation of the WT and WT control parabiosis, at 8-9 weeks. D. Frequency of wild-type (WT) B cells in the circulation of 564Igi parabiosis partner (grey circles) or WT parabiosis partner (control parabiosis, black circles) over time. Week 1 (564Igi n=11 vs WT n=6), Week 2 (n=5 vs n=6), Week 3 (n=10 vs n6), Week 4 (n=8 vs n=4), Week 5 (n=10 vs n=6), Week 6 (n=10 v n=6), Week 7 (n=9 vs n=2), Week 8-9 (n=10 vs n=6). E. Frequency of 564Igi (blue circles) or WT (control parabiosis, yellow circles) B cells in the circulation of WT parabiosis partner mouse over time. As in Figure 1B, but week 2 (564Igi n=6 vs WT n=6), Week 6 (n=9 vs n=6).

F. Frequency of 564Igi B cells in WT parabiosis partner (left column), and in the 564Igi partner, the proportion of WT B cells in the total splenic B cell population or within the germinal center (GC) B cell population within the spleen 8 weeks following surgery.

G. Parabiosis pairing of Toll-Like receptor 7 knock-out (TLR KO) mouse with 564Igi mouse for 8 weeks, showing the proportion of TLR7 KO B cells within circulation (n=5), splenic B cell pool (n=6) and splenic GC B cell population (n=6) of the 564Igi parabiosis partner.

H. Parabiosis pairing of Toll-Like receptor 7 knock-out (TLR KO) mouse with 564Igi mouse for 8 weeks, showing the proportion of TLR7 KO B cells within the mesenteric lymph node (mesLN) B cell pool (n=6) and mesLN GC B cell population (n=6) of the 564Igi parabiosis partner.

Significance is indicated as ***P<0.001, **P<0.01, *P<0.05, and P>0.05 (not significant (NS)) by Mann-Whitney test for Fig 1D, 1E and by Wilcoxon paired test for Fig 1F, 1G, 1H.

To address whether the increased competitiveness of WT B cells in 564Igi parabionts was due to the relative abundance of B cells in WT mice, we performed WT parabiosis with 564Igi heterozygous mice (50% WT, 50% 564Igi B cells), which have at least 2-fold more B cells than 564Igi homozygous mice. However, the circulatory B cell exchange kinetics in parabionts of WT mice with either 564Igi heterozygous or 564Igi homozygous mice were comparable (Suppl.Fig.1C-F), indicating that it was not the increased number of B cells that made the WT cells more competitive. WT B cell dominance of the splenic GCs in the 564Igi parabiont was also dependent on Toll-like receptor 7 (TLR7) expression (Fig.1G), consistent with our earlier observations of TLR7-dependence in WT B cell participation in the GC response in the 564Igi:WT bone marrow chimera system (*7*). Consistently, in the mesenteric lymph node (mesLN), which is primarily targeted at microbial antigens, WT B cell GC residence was TLR7 independent (Fig.1H). We next assessed the reactivity of WT-derived serum antibodies to nucleolar autoantigens.

Due to allotype differences in the heavy chain locus of 564Igi and WT B cells, the source of IgG2 antibodies can be distinguished by two isoforms, i.e. IgG2a (564Igi) and IgG2c (WT) respectively. We found increased reactivity of IgG2c antibodies against nucleolar auto-antigens in the sera of 564Igi:WT parabionts in contrast to control parabiosis (WT:WT)(Suppl.Fig.1G). Together these results mirror the findings from the earlier mixed bone marrow chimera model, where WT B cells outcompete the spontaneous GC-initiating 564Igi clone and generate autoantibodies but with the advantage to observe the WT B cell ability to dominate mature GCs in the 564Igi mice.

### Enrichment of WT B cells in GCs is persistent

To examine the dynamics of WT B cell invasion and their persistence in autoreactive GCs without a continuous WT B cell source, parabiotic mice were separated at 2 weeks following surgery (Fig.2A). This time point was chosen as this is when parabionts reached adequate cross-circulation, as indicated by T cell equilibration. Following separation, the abundance of WT B and T cells gradually decreased in the circulation of 564Igi recipients, displaying a similar trend to control separation parabionts (Fig.2B,C, Suppl.Fig.2A,B). Of note, WT B cells in the 564Igi parabiont only constituted approximately 20% of peripheral blood B cells 6-7 weeks following separation. By contrast, in the spleen, WT B cells were dominant. Specifically, WT B cells constituted more than 80% of all splenic GC B cells 6-7 weeks after separation in the 564Igi mouse (Fig.2D). WT B cells also dominated the GC response in the axillary/brachial and mesenteric LNs (Suppl.Fig.2C). Importantly, splenic GC enrichment was not apparent at the time of parabiotic separation (Suppl.Fig.2D) but was present for up to a year post-separation (Suppl.Fig.2E). Together, these results show that WT B cells can invade and persist in the autoreactive GC without a continuous source.

**Figure 2.**
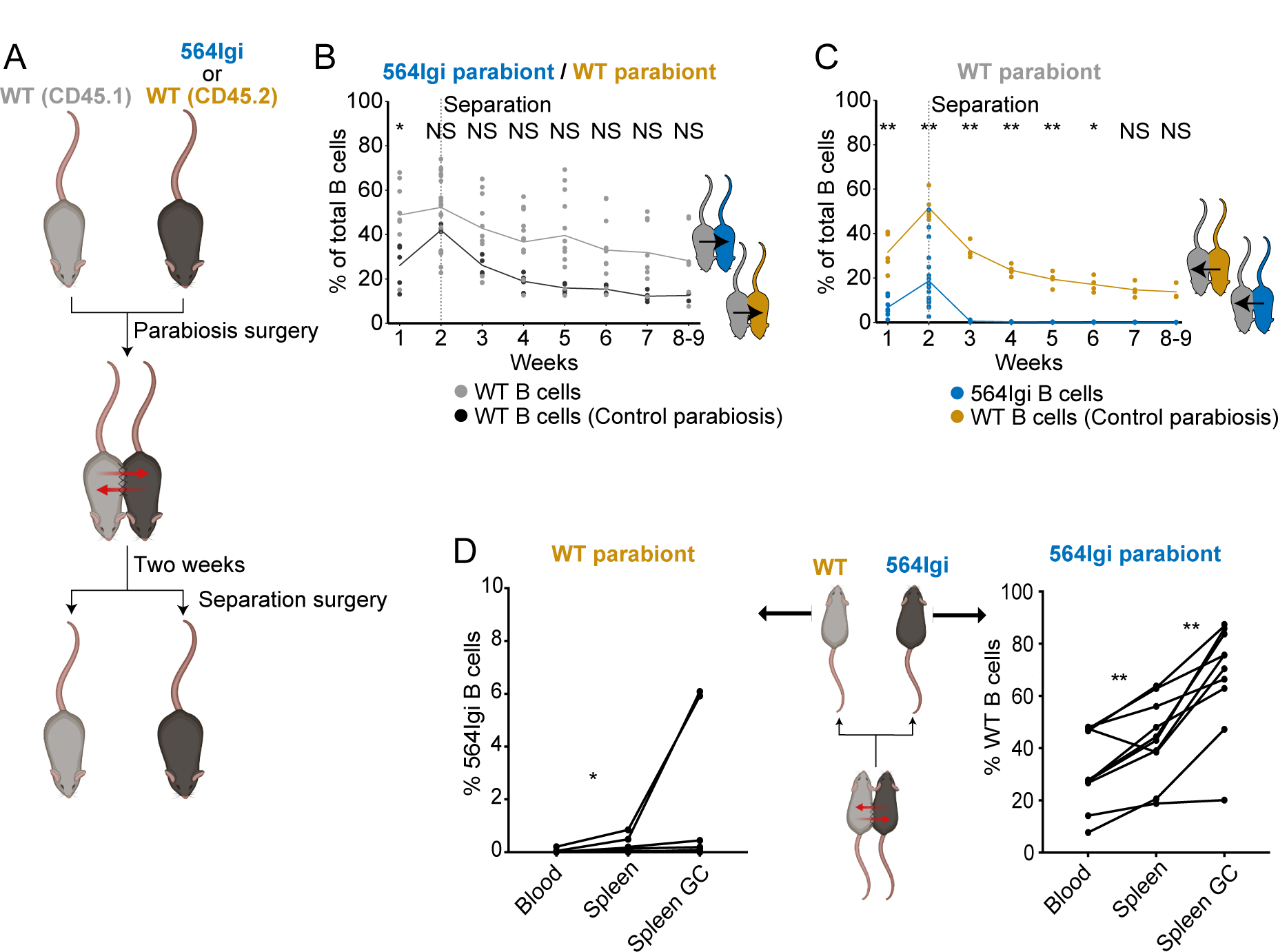
Temporary parabiosis highlights WT B cell persistence in the spontaneous germinal center. A. Schematic representation of parabiosis surgery followed by surgical separation at 2 weeks of parabiosis and subsequent follow-up. B. The proportion of 564Igi (blue circles) or WT (Control parabiosis, yellow circles) B cells in the circulation of WT parabiosis partner mouse over time. Week 1 (564Igi n=8 vs WT n=5), Week 2 (n=19 vs n=5), Week 3 (n=12 vs n=4), Week 4 (n=11 vs n=4), Week 5 (n=11 vs n=4), Week 6 (n=9 v n=4), Week 7 (n=7 vs n=4), Week 8-9 (n=8 vs n=3). C. The proportion of wild-type (WT) B cells in the circulation of 564Igi (grey circles) or WT (control parabiosis, black circles) parabiosis partner mouse over time. Number of mice as in Figure 2B. D. The proportion of 564Igi B cells in the WT parabiosis partner (left graph, n=9), and in the 564Igi parabiosis partner (right graph, n=10), the proportion of WT B cells in the circulation, splenic B cell population or splenic germinal center B cell population, after 2 weeks of parabiosis and subsequent 6 weeks of separation. Significance is indicated as ***P<0.001, **P<0.01, *P<0.05, and P>0.05 (not significant (NS) by Mann-Whitney test for Fig 2B, 2C and Wilcoxon paired test for Fig 2D (Blood vs Spleen, Spleen vs Spleen GC).

### WT B cells clonally expand in spontaneous GCs

To determine if the WT B cells underwent clonal selection and expansion in 564Igi GCs, Aicda-Cre^ERT2^ Confetti reporter mice were attached to 564Igi mice for two weeks, then they were separated and the 564Igi ex-parabiont was treated with tamoxifen. Upon tamoxifen induction, Aid-expressing (GC) B cells and their progeny express 1 of 10 different color combinations (*9*), allowing us to visualize the clonal dynamics of these B cells in the spontaneous GCs (*9–11*). At several time points post-separation, spleens were harvested, and their sections analyzed for Confetti-expressing B cells within GCs (Fig.3A). The possibilities are that, upon activation and AID expression, (1) many individual B cells of different clonalities are equally capable of expanding, and as such, they undergo minimal clonal selection and maintain the original color diversity; (2) fate-mapped B cells do not persist in the GC and are replaced by newly recruited B cells that do not show any of the Confetti colors (dark clones), thus diminishing fate-mapped B cells over time; or (3) specific clones are selected and expand, giving rise to the dominance of a single or few colors (referred to as pauci-clonality). The latter is observed in foreign antigen responses and also in 564Igi:WT bone marrow chimeras (*7, 9*). Indeed, we also observed that fate-mapped B cells in the GCs of 564Igi parabionts gradually lost color diversity, indicating active participation and clonal selection of these B cells in the spontaneous GCs (Fig.3B-D). Despite the overall proportion of WT B cells increasing over time after separation in our experiment above (Fig.2D, Suppl.Fig.2D), the total number of fate-mapped B cells per GC decreased from day 14 to 30 post-tamoxifen (Fig.3E), suggesting that non-fate-mapped cells (dark clones), likely naïve B cells, are the source of this B cell enrichment. Together, these results show that after a transient WT B cell source, WT B cells clonally expand in the autoreactive GC and contribute, in part, to WT B cell enrichment in the GC.

**Figure 3.**
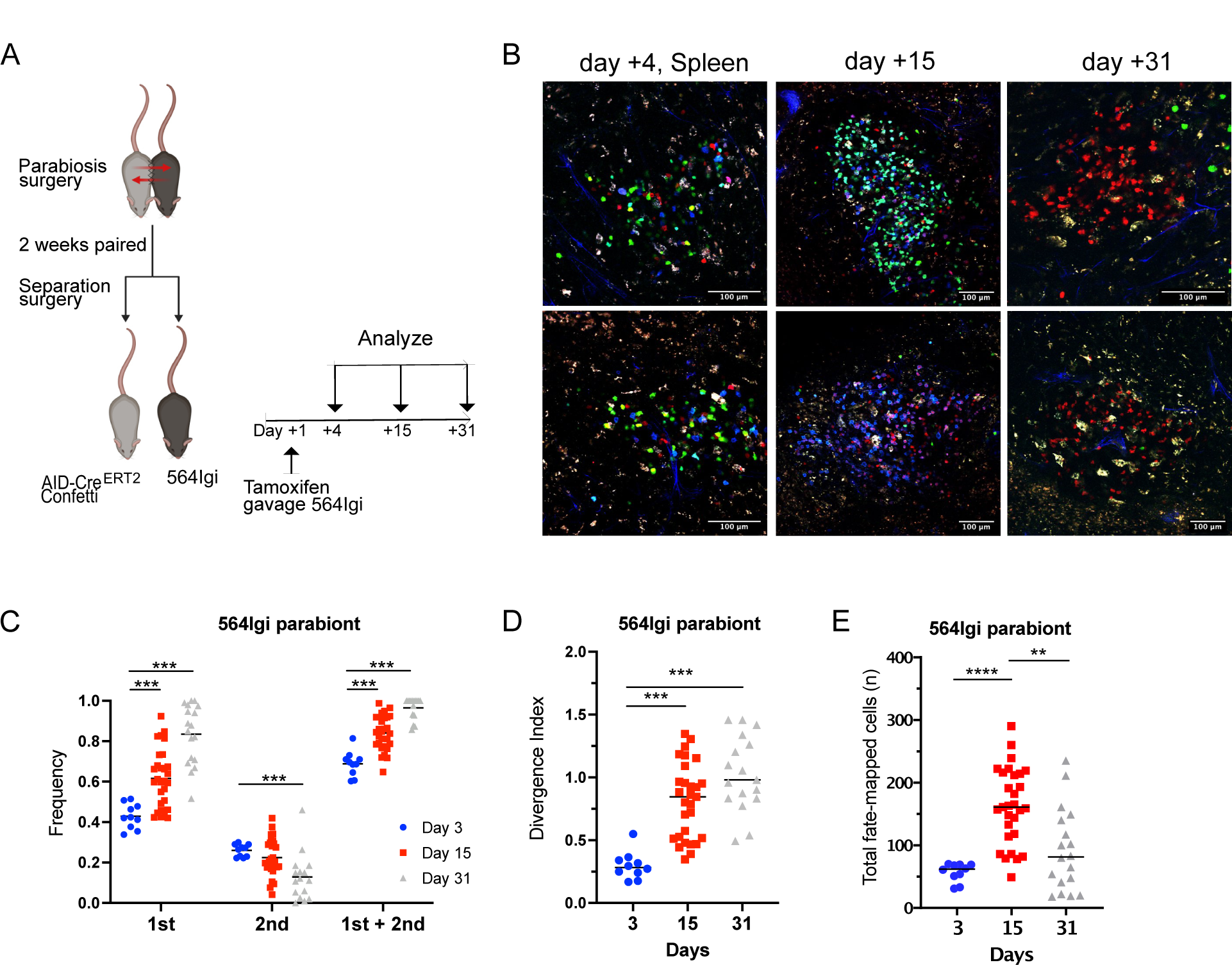
Temporary parabiosis highlights WT B cell persistence and clonal expansion in autoreactive 564Igi GCs. A. Schematic depiction of parabiosis and separation surgery of AID-Cre^ert2^ Confetti mouse with 564Igi mouse to assess clonal evolution of WT B cells in 564Igi partner at day +4, +15, and +31 post-separation surgery. B. Representative multi-photon images of the splenic GC of the 564Igi parabiosis mouse partner for quantification of color dominance. The scale bars indicate 100 µm. C. Frequency of most frequent color (1st), second-most frequent color (2nd), and their sum observed in individual splenic GC at Day 3, 15, and 31 post-tamoxifen induction of Confetti recombination in 564Igi parabiont. Day 3 (2 mice, 10 GC), day 15 (3 mice, 30 GC) and day 30 (4 mice, 18 GC). D. The divergence index for the GC represented in (C). E. The number of fate-mapped WT B cells for the GC represented in (C). Significance is indicated as ***P<0.001, **P<0.01, *P<0.05, and P>0.05 (not significant (NS)) by Mann-Whitney test for Fig 3C,3D (compared to day 3) and ANOVA with posthoc test (Dunn) for Fig 3E.

### WT B cells enter the pre-existing autoreactive GCs within days

Having observed the engraftment of WT B cells in the autoimmune environment as early as 2 weeks following parabiosis surgery, an adoptive transfer approach was established to further characterize and identify the kinetics of WT B cell participation in autoreactive GCs. WT B cells were adoptively transferred to 564Igi recipients, and spleens were harvested at varying periods post-transfer (Fig.4A). As early as day 4 post-adoptive transfer, WT B cells participated in the GC response. By day 7 post-transfer, the proportion of transferred WT B cells within the total GC B cell pool was significantly increased over the proportion of WT B cells within the total B cell compartment of the spleen (Fig.4B,C). Early enrichment of donor B cells was specific to the autoreactive GC, as it was not seen in the mesenteric lymph node and Peyer’s patches of the 564Igi recipient or in immunized WT mice (Suppl.Fig.3A-D). Of note, we saw little engraftment of WT B cells transferred to unimmunized WT hosts (Suppl.Fig.3A-D). WT B cells were frequently found in 564Igi GCs, with most GCs in the 564Igi recipient mice populated by WT B cells (Fig.4D). This was consistent among 564Igi recipients despite differences in the number of white pulp areas and GCs (Fig.4E). A range of WT B cell residency patterns within GCs – from dominant to sparse – was observed (Fig.4F), but there were no GCs composed solely of donor B cells at day 7 post-adoptive transfer. Thus, we conclude that, within four days post-transfer, donor B cells joined existing GCs. WT B cell entry of GCs may be dependent on the size of the GC. Conversely, the size of the GC could be impacted by WT B cell residency. However, the GC size was comparable between GCs with both donor and recipient residents and GCs with 564Igi B cells only (Fig.4G). Together, these data show that, like foreign-antigen-driven GCs, autoreactive GCs are open structures to which B cells can be recruited rapidly (*12*).

**Figure 4.**
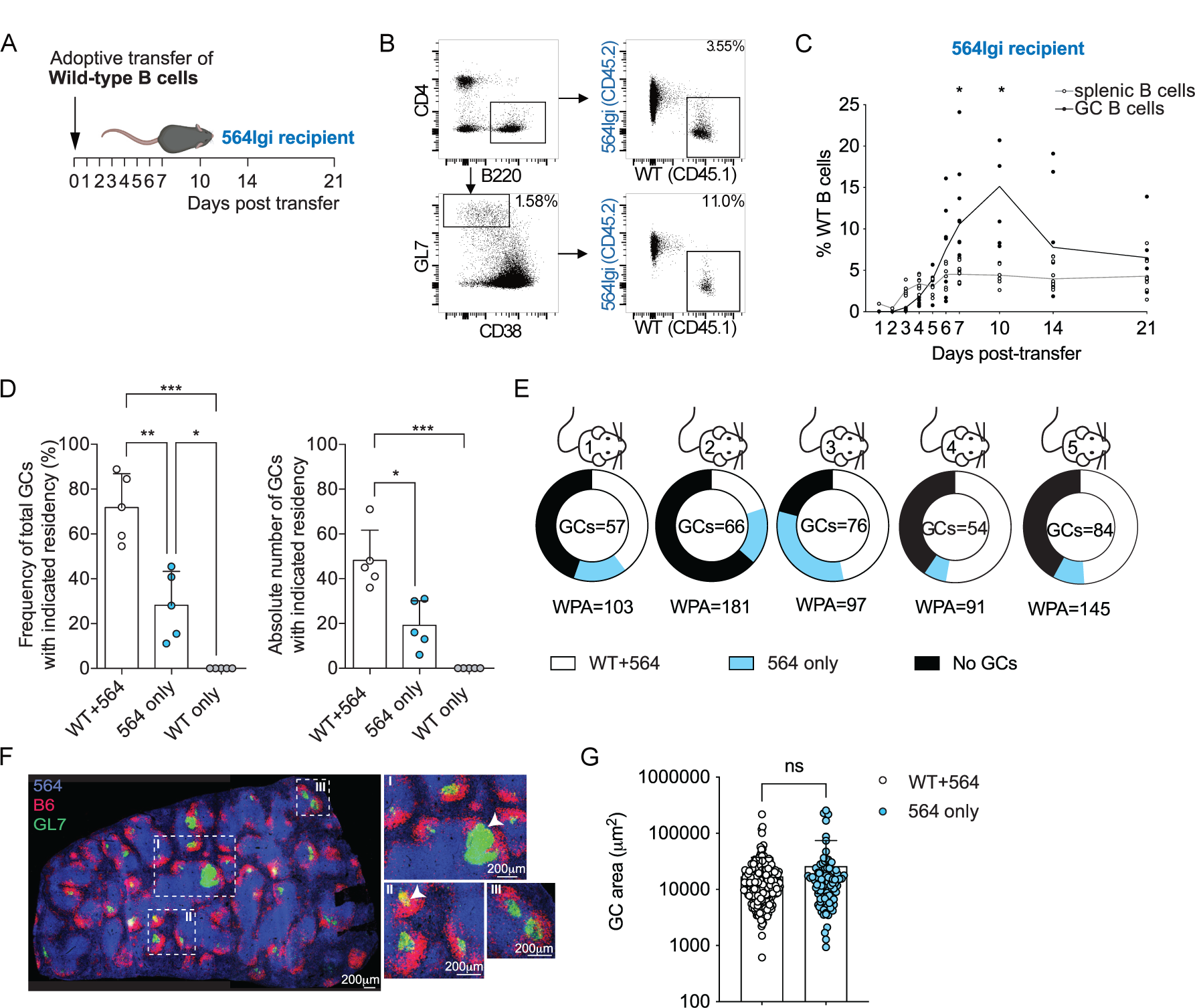
Wild-type B cells rapidly enter the pre-existing autoreactive germinal center. A. Schematic depiction of WT B cell adoptive transfer model and follow-up time points. B. Representative FACs dot plot of WT B cells in the splenic B cell compartment and within the splenic germinal center B cell compartment, at day 7. Day 1 (n=1), day 2 (n=1), day 3 (n=7), day 4 (n=7), day 5 (n=7), day 6 (n=7), day 7 (n=11), day 10 (n=6), day 14 (n=9), day 21 (n=7), day 56 (n=5). C. The proportion of WT B cells within the splenic B cell pool (grey line, open circle) and splenic germinal center B cell pool (black line, closed circle) over time. D. Bar charts indicating the frequency (left) and absolute number (right) of GCs populated by WT and 564Igi, 564Igi only, or WT only B cells (left). Each symbol indicates a mouse. E. Pie charts demonstrating the frequency of GCs populated by both WT and 564Igi, 564Igi only or WT only B cells as a frequency of white pulp areas (WPAs) in individual mice. No GCs with only B6.2 B cells were identified. F. Representative confocal immunofluorescence microscopy image showing GC composition in a splenic section; GCs were identified by GL7 (green) staining and confirmed by the overlapping CD21bright staining of FDC networks in a consecutive slice. Anti-CD45.1 (blue) and anti-CD45.2 (red) staining was used to identify 564Igi-and WT-derived cells, respectively. Arrows indicate GCs with different patterns of WT residency. GCs contained within boxes with roman numerals are magnified in the smaller panels to the right. G. Bar chart showing the area of GCs with WT and 564Igi and 564Igi only residents. Each symbol indicates a GC. Wilcoxon paired test: Fig 4B. Fig 4C: One-way ANOVA with Bonferroni’s multiple comparison test. Fig 4F: Mann-Whitney test. Values are mean+/-SD. Significance is indicated as ***P<0.001, **P<0.01, *P<0.05, and P>0.05 (not significant (NS)).

### WT B cells actively participate in autoreactive GCs and can differentiate into autoantibody-producing cells

We next aimed to confirm that adoptively transferred WT B cells actively engaged in the GC response in contrast to transient occupancy (*13*). To address this, we used B cells from S1PR2-Cre^ERT2^TdTomato reporter animals; TAM administration permanently labels *S1pr2*-expressing GC and activated B cells and their progeny with tdTomato (*14*). We transferred S1pr2 reporter B cells to 564Igi Aicda-Cre^ERT2^EYFP recipients. Notably, S1pr2-TdTomato-expressing donor B cells (WT) co-localized in 564Igi GCs with Aicda-Cre^ERT2^EYFP-expressing 564Igi B cells (Suppl.Fig.3E). Interestingly, donor-derived antibodies (IgG2c) specific for nuclear antigens (ANA), neutrophil cytoplasmic antigens (ANCA) and ssDNA were present as early as 7 days post-adoptive transfer (Suppl.Fig.3F-H). After transfer, the relatively quick rise of WT-derived plasma cells before the peak of donor GC B cells suggests that both extra-follicular and GC-derived antibody-producing plasma cells are generated (Suppl.Fig.3I).

To demonstrate that GC-derived B cells possessed the ability to produce autoantibody at this early time-point, S1pr2-TdTomato reporter B cells (WT B cell repertoire) were transferred into tamoxifen-treated 564Igi recipients. TdTomato-expressing (S1pr2^+^) B cells were then sorted and cultured, then the antibody reactivity of GC-experienced WT B cells was measured in the cell culture supernatant (Suppl.Fig.3J). Self-reactive antibody against ssDNA was detected in sorted TdTomato-expressing (S1pr2^+^) B cells. We further observed that sorted donor B cells that did not express the S1pr2 reporter also secreted ssDNA antibody, although at a lower level, suggesting a GC-independent source of auto-antibody production or incomplete labeling of B cells in the GC in this narrow timeframe of tamoxifen exposure (Suppl.Fig.3K). Together, these results show that donor-derived WT B cells can enter the autoreactive GC and subsequently differentiate into autoantibody-producing cells.

### GC invasion by WT B cells is dependent on BCR specificity and MHC expression and can be regulated by TLR7 and IFNAR expression

The adoptive transfer approach allowed us to further examine the molecular requirements for B cell entry into spontaneous GCs and to identify possible targets for intervention. The role of BCR specificity for entry into GC was assessed by the transfer of B1-8 transgenic (Tg) B cells into 564Igi recipients. In the B1-8 Tg mouse strain, the pairing of the Ig heavy chain knock-in with Ig lambda (Igλ) light chain confers specificity to the hapten 4-hydroxy-3-nitrophenylacetate (NP); thus approximately 5% of the peripheral B cells are NP-specific (Fig.5A) (*15*). In contrast to observations with WT B cells, the proportion of Igλ^+^ Tg B cells (NP-specific) within the GC population was decreased relative to naïve B1-8-derived B cells in both spleen and mesLN (Fig.5B, Suppl.Fig.4A). Thus, NP-specific B cells did not participate in the spontaneous GC response, indicating the essential role of antigen recognition in this process. NP-specific Igλ+ B cells did however readily enter GCs in WT mice immunized with haptenated NP-CGG in a similar adoptive transfer scheme (Suppl.Fig.4B), as previously published (*16*). In contrast to the results with NP-specific B cells, we found that Sle1yaa-derived B cells not only readily enter the GC when transferred into 564Igi recipients, but their capacity to invade GCs was superior to the B cells derived from WT mice (Fig.5C). Sle1yaa mice have the lupus erythematosus susceptibility locus as well as a Y chromosome translocation that increases the expression of TLR7 approximately 2-fold, which suggests a role for increased TLR7 expression in enhancing GC entry (*17*).

**Figure 5.**
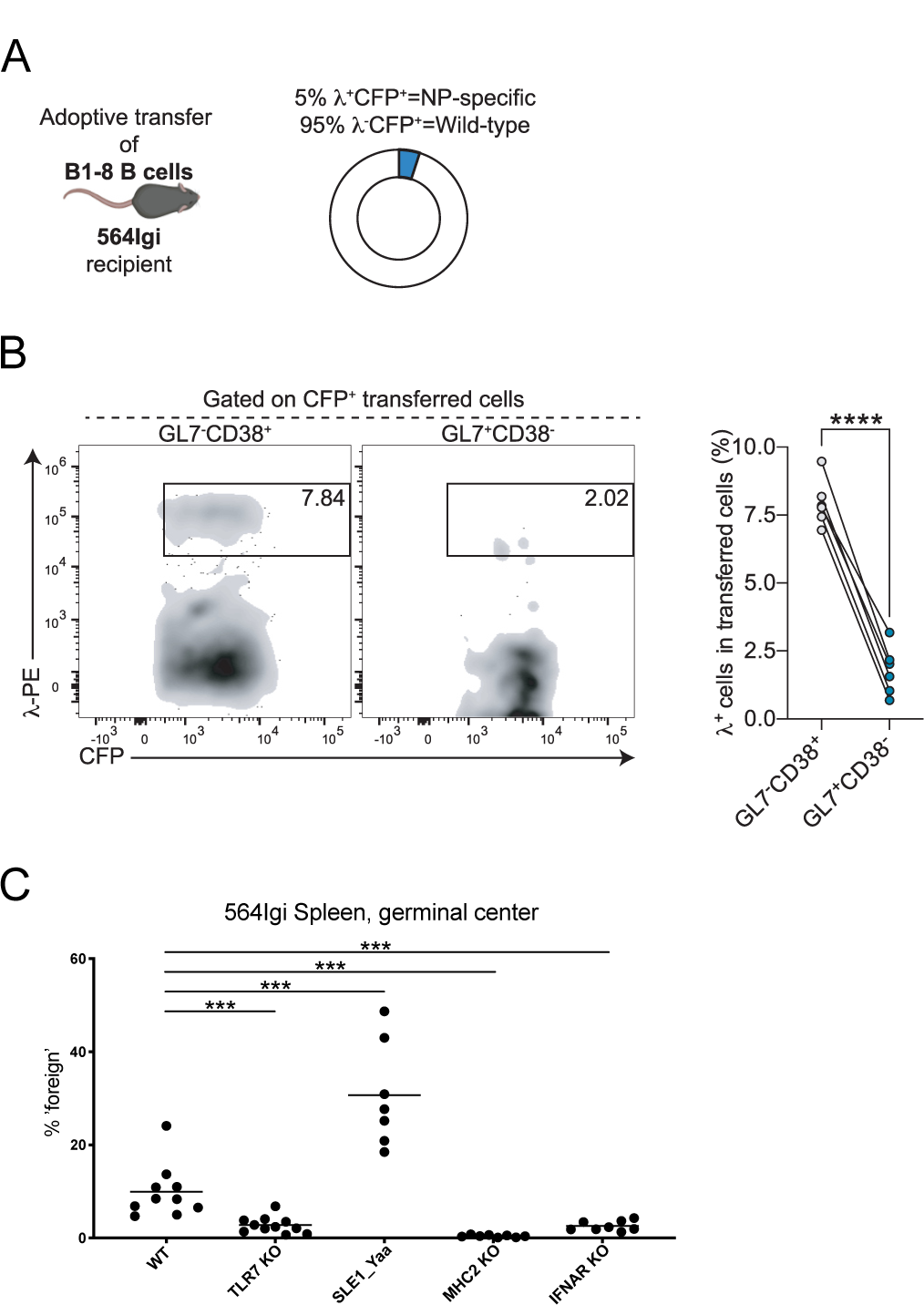
Molecular requirements for wild-type B cell entry and participation in spontaneous germinal centers. A. Schematic representation of B1-8 B cell adoptive transfer. B. Representative FACSplot and gating for identifying NP-specific B cells (left panel). The proportion of Lambda-positive (NP-specific) B cells within the splenic B cell compartment, splenic GC B cells in the 564Igi mouse, 7 days post-transfer (n=6) (right panel). C. The proportion of germinal B cells derived from WT, MHC2 KO, TLR7 KO, SLE1-Yaa, and IFNAR KO transferred B cells within the splenic GC B cell compartment, 7 days post-transfer. WT transfer n=10, MHC2 KO transfer n=8, TLR7 KO transfer n=11, SLE1-Yaa n=7, IFNAR KO transfer n=8. Significance is indicated as ***P<0.001, **P<0.01, *P<0.05, and P>0.05 (not significant (NS)). The statistical test used is the Mann-Whitney test for Fig 5B, C (compared to WT).

To determine the requirement for antigen presentation in the participation of WT B cells in spontaneous GCs, MHCII KO B cells were adoptively transferred into 564Igi recipients (Fig.5C). Notably, MHC II–deficient B cells failed to participate in spontaneous GCs, even when MHCII expression was halved (MHCII haploinsufficiency) (Suppl.Fig.4C). MHCII–deficient B cells were able to enter the B cell follicle but not the GC area (Suppl. Fig.4D). B cells similarly failed to engage in the GC response upon adoptive transfer when they lacked the expression of TLR7 or type I interferon receptor (IFNAR) signaling (Fig.5C). Diminished expression of IFNAR (IFNAR haploinsufficiency) did, however, result in partial recovery of B cell participation in the GC response (Suppl.Fig.4C). Of note, both TLR7-KO and IFNAR-deficient B cells competed similar to WT B cells for entry into the GC of the mesLN (Suppl.Fig.4E), suggesting selective TLR7 and IFNAR dependence towards the autoreactive GCs. Together, these data demonstrate that WT B cell entry and participation in spontaneous GCs is an active process dependent on BCR specificity and antigen presentation and can be titrated by the expression level of TLR7 and IFNAR.

## DISCUSSION

In a foreign antigen response, established germinal centers are dynamic, open structures that harbor B cells with different specificities and allow naïve B cells to enter and participate in the GC process (*12, 15, 16, 18–22*). We found that, within the 564Igi mice, WT B cells were able to enter and participate in the autoimmune GC response, eventually acquiring a specificity for ssDNA and also displaying a wide array of class-switched autoantibody reactivities, suggesting possible recruitment of B cells with other specificities. Our observations are reminiscent of the breach of B cell tolerance observed in tonsil biopsies of SLE patients, where despite autoreactive B cells being excluded from the GCs of healthy individuals, in SLE patients, autoreactive B cells readily engaged in the GC response (*23*).

Progressive GC enrichment of WT B cells is evident in the parabiosis model, yet there was a decrease in the number of fate-mapped B cells after temporary parabiosis. This suggests that WT naive B cells (dark cells) entered the GC, as is observed in long-lasting GCs primed by respiratory viruses (*20*). In the adoptive transfer model, from 10 days post-transfer onwards, the proportion of WT B cells decreased and persisted at a lower level, supporting that persistent WT B cell GC enrichment may be dependent on a continuous source of naïve B cells. WT B cells did increasingly invade the GCs of the mesenteric lymph node (mesLN) and Peyer’s patches (PP) and persisted at 8 weeks post-transfer. This could suggest that WT B cells initially favor entry into the autoreactive GC, but at a later stage, the majority of naïve B cells preferentially enter and persist in the mesLN and PP, as they are likely more reactive than 564Igi B cells to the microbial antigens found in gut-related lymph nodes.

In murine infection and vaccination response models, the entry of naive B cells into GCs depends on specific BCR and MHCII expression, as in our autoimmune models (*12, 16, 20, 22, 24, 25*). In contrast, we show that TLR7 and IFNAR expression altered the threshold for naive B cell entry in the autoreactive GC but minimally affected entry in the mesenteric lymph node. This difference is of interest as it could potentially be utilized to specifically address the autoreactive GC process, thus could aid in the identification of early intervention strategies that could halt or slow disease progression. TLR7 signaling is important in the establishment of autoimmune diseases, such as SLE, by inducing extrafollicular antibody-forming cells (*26, 27*). The dual function of TLR7 might highlight the requirements for the initiation of disease by autoreactive B cells (GC-independent) and, at least in part, the propagation of disease by recruitment of new B cells (GC-dependent). IFNAR signals play an important role in the regulation of TLR7 responses in B cells and could contribute to further GC-dependent propagation of disease, as B cell–intrinsic type 1 IFN receptor signals are not required for SLE development but do enhance disease (*28–30*). Lack of GC-specific IFNAR signaling indeed reduces the size and frequency of the GC as well as auto-antibody production, and our data suggest that this might be due to reduced B cell entry to the GC (*31, 32*).

Our findings not only give mechanistic insight into the kinetics and requirements for a break of tolerance of naïve B cells but also underscore the potential therapeutic targets in autoimmunity. The chronic and progressive nature of WT B cells joining the GC response calls for early interventions. Total blockade of GC responses might, however, exacerbate autoimmune disease (*33, 34*), and warrants selective GC checkpoints such as TLR7 and IFNAR to regulate autoreactive from foreign GC responses, but could conflict with responses to viral infection (*35, 36*). We suggest that adoptive transfer in 564Igi mice can be utilized more widely as a novel tool to address the early events and molecular mechanisms facilitating the break in tolerance of naive B cells.

## MATERIALS AND METHODS

### Study design

We performed parabiosis and adoptive transfers to assess the kinetics of WT B cell participation and break of tolerance in the autoreactive GC. The details of the biological replicates in each group and the number of repetitions per experiment are indicated in the respective figure legends. Experimental end-points were established before any *in vivo* experiments for animal welfare considerations. All animals used for comparison within each experiment were age- and sex-matched. Mice in different groups were randomly assigned, outliers were not excluded, and studies were unblinded.

### Mice

This study was conducted in accordance with the NIH Guide for the Care and Use of Laboratory Animals. Animal experiments were performed as specified in protocols approved by the local IACUC of Harvard Medical School (protocol number IS00000111). All surgeries and cell transfers were performed under isoflurane anesthesia, and every effort was made to minimize suffering. 564Igi mice were provided by Theresa Imanishi-Kari (Tufts University) and maintained in-house, and different congenic backgrounds (CD45.1 and CD45.1.2) and a 564Igi AicdaCreERT2-YFP were generated. AicdaCreERT2-Confetti, B1-8, and S1PR2CreERT2-TdTomato donors were provided by Gabriel D. Victora (Rockefeller University). B6 (C57BL/6J), B6 CD45.1 (B6.SJL-Ptprca Pepcb/BoyJ), TLR7 KO (B6.129S1-Tlr7 tm1Flv /J), I-AB-flox (MHCII-fl) (B6.129X1-H2-Ab1tm1Koni/J), Sle1Yaa (B6.Cg-Sle1NZM2410/Aeg Yaa/DcrJ), MB1-Cre (B6.C(Cg)-Cd79atm1(cre)Reth/EhobJ) were from Jackson Laboratories and maintained in house. IFNaR fl/fl (Ifnar1tm1Uka) mice were provided by Ulrich Kalinke. MB1-Cre mice were crossed with I-AB-flox and with IFNaR fl/fl to generate B cell-specific expression. AicdaCreERT2 flox-stop-flox-EYFP mice were from Claude-Agnes Reynaud and Jean-Claude Weill (Institut Necker) and crossed with 564Igi mice. To generate 564IgiHet, MHCII+/-, and IFNaR +/-mice, corresponding homozygous mice were crossed with B6, B6 CD45.1, or MB1-Cre and maintained in-house.

| Experimental models: Organisms/strains | Source | Identifier |
| --- | --- | --- |
| Aid-CreERT2 EYFP: AicdaCreERT2 flox-stop-flox-EYFP | Dogan et al., 2009 | Ref (13) |
| 564Igi | Berland et al., 2006 | Ref (8) |
| Aid-CreERT2 Confetti: AicdaCreERT2-Rosa26Confetti | Tas et al, 2016 | Ref (9) |
| C57BL/6j (CD45.2) | The Jackson Laboratory | 000664 |
| B1-8 (PA-GFP) | Victora et al, 2010 | Ref (37) |
| S1PR2-CreERT2-TdTomato | Shinnakasu et al., 2016 | Ref (14) |
| B6.SJL-Ptprca Pepcb/BoyJ (CD45.1) | The Jackson Laboratory | 002014 |
| TLR7 KO: B6.129S1-Tlr7 tm1Flv /J | The Jackson Laboratory | 008380 |
| I-AB-flox: B6.129X1-H2-Ab1tm1Koni/J | The Jackson Laboratory | 013181 |
| Sle1Yaa: B6.Cg-Sle1NZM2410/Aeg Yaa/DcrJ | The Jackson Laboratory | 021569 |
| MB1-Cre: B6.C(Cg)-Cd79atm1(cre)Reth/EhobJ | The Jackson Laboratory | 020505 |
| Ifnar1 tm1Uka | Le Bon et al. 2006 | Ref (38) |

Mice were bred and maintained in an AAALAC-accredited facility at Harvard Medical School under specific pathogen-free (SPF) conditions. Both female and male mice were used in the adoptive transfer studies. Only females are suited for parabiosis experiments. Mice were on a standard 12hr light/dark cycle and were kept on a high-fat diet for 1-2 weeks before and during the parabiosis experiments and on a regular low-fat diet for adoptive transfer studies. Mice were 7–9 weeks old at the start of experiments. Mice were age-matched when comparing different strains (max 1 week between experimental and control groups).

### Antibodies and staining reagents

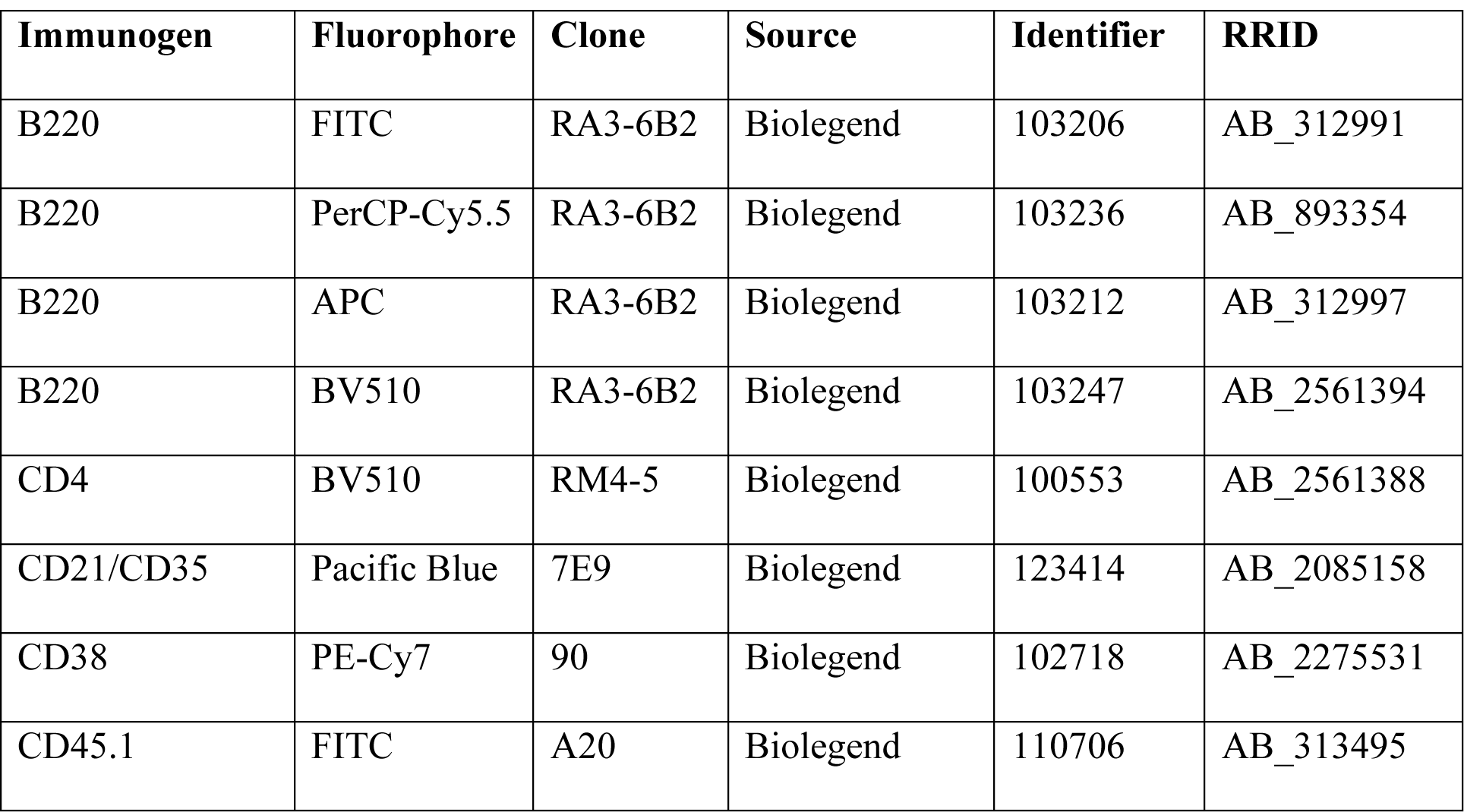
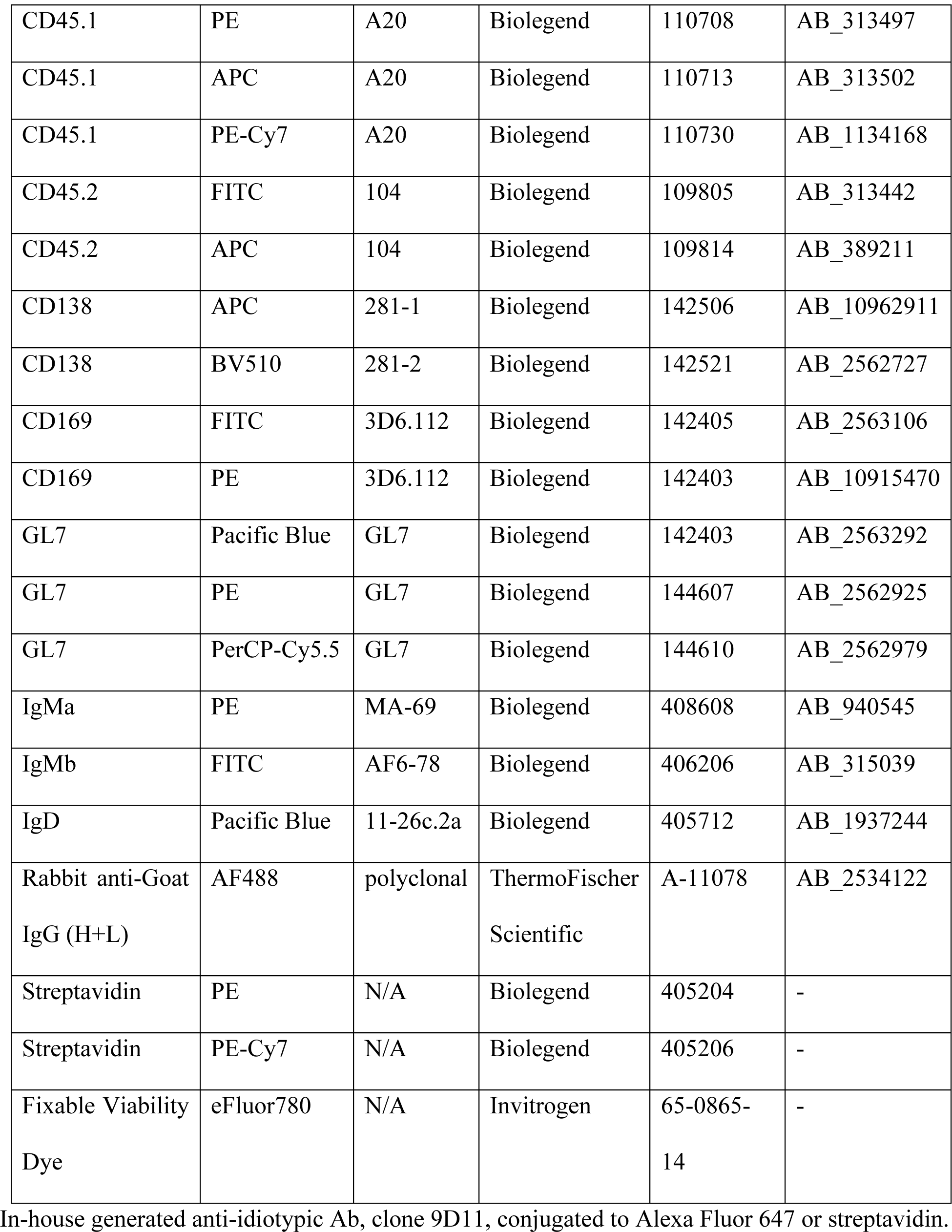

### Genotyping and FACS-typing mice

Genotyping was performed using the primers and reaction conditions indicated in Table S1. For FACStyping, 564Igi mice were bled retro-orbitally. Using heparinized capillary tubes, approximately 60–100 µl blood was drawn into Eppendorf tubes containing 30 µl acid-citrate-dextrose solution. Following collection, the stabilized blood was briefly spun down, then under-layered with 1 ml of Lymphocyte Separation Medium, and spun for 25 min at 1,600 RPM at RT. The mononuclear cell layer was aspirated and transferred into 1 ml ice-cold FACS buffer, mixed, then pelleted at 200 g for 5 min. Cells were resuspended in FACSbuffer and processed for flow cytometric analysis as described further below. FACStyping of 564Igi mice was performed using B220, anti-IgMa, anti-IgMb, and 9D11. 564Igi mice were genotyped by ddPCR using primers (Table S1) to assess the expression of heavy and light chains.

### Parabiosis surgery

Mice were co-housed for 1-2 weeks before parabiosis surgery, and their flanks were shaved a day before. Surgeries were performed on heating pads under aseptic conditions with controlled isoflurane anesthesia. Mirror-image incisions at the left and right flanks of the mouse were made through the skin. Elbow and knee joints from each parabiont were sutured together (Ethilon 4-0), followed by a continuous suture to join the skin together (Ethilon 4-0). Mice in each pair were injected subcutaneously with buprenorphine twice a day for 3 days or buprenorphine SR once with daily meloxicam to manage pain, supplemented with 0.9% (w/v) sodium chloride for hydration. Control parabiosis surgeries consisted of joining wild-type mice differing in congenic marker (CD45.1 and CD45.2). In addition, the side of the surgery was alternated between similar parabiosis pairings. Mice were housed in alpha-dry bedding and received sulfamethoxazole/trimethoprim in their drinking water for 10 days. For overall health and maintenance behavior, mice were monitored daily for the first week and analyzed for weight loss, grooming, stress, and pain response, followed by visual monitoring 2-3 times a week.

The parabiosis separation surgery was performed after 2 weeks of pairing and followed the same setup, analgesia, and monitoring as the initial parabiosis surgery. The sutures joining the knee and elbow joints were removed and the skin of each mouse was closed with a continuous suture (Ethilon 4-0). For blood collection mice were bled retro-orbitally and via the submandibular route. Using heparinized capillary tubes, approximately 60–100 µl blood was drawn into Eppendorf tubes containing 30 µl acid-citrate-dextrose solution.

### B cell isolation and adoptive transfer

Spleens were passed through a 70-um cell strainer, and splenocytes were resuspended in RBC lysis buffer (155 mM NH4Cl, 12 mM NaHCO3, 0.1 mM EDTA), incubated for 2–3 minutes then spun down as before and resuspended in MACS buffer. Splenocytes were incubated with 100ul of biotin-antibody cocktail for 1×10^8 cells in 1ml of MACS buffer (2% fetal calf serum, 1mM EDTA) for 15min on ice, followed by 100ul of Streptavidin nanobeads for 1×10^8 cells in 1ml of MACS buffer for 15min on ice (MojoSort Mouse pan B cell isolation kit II, Cat.no. 480088, Biolegend). The cell suspension was diluted and put on a pre-rinsed LS Column (Cat. No. 130-042-401, Milteny Biotec), and the column was washed three times. The flow-through was collected on ice, counted and diluted until 10×10^6 cells per 100ul of HBSS and kept on ice. Purity was assessed by FACS and showed consistent >96% B cell (B220+) purity. The adoptive transfer was performed following appropriate dilution by retroorbital injection on anesthetized mice.

Adoptive transfer following immunization was done after 7 days of NP-CGG immunization and 11 days after the first SRBC immunization. For the adoptive transfer of NP-specific B cells, mice were immunized with 100ug of NP(24)-CGG (Cat. N5055C-5, Biosearch technologies) emulsified in alum (Cat. 77161, Thermo Fisher Scientific). The following day, isolated B cells from B1-8 mice were adoptively transferred (10*10^^^6 B cells/recipient); mice were sacrificed on day 7 post-transfer.

564Igi recipients were placed on a tamoxifen diet (TD130860, Envigo) for 3 weeks before B cell transfer and maintained on a tamoxifen diet until the end of the experiment. B cells were isolated from S1pr2-TdTomato reporter mice and adoptively transferred to 564Igi animals (10*10^^^6 B cells/recipient). On day 7 following adoptive transfer, 564Igi recipient animals were sacrificed. Following RBC lysis, splenocytes were stained with antibodies against IgD, CD138, B220, GL7, CD45.1, and viability dye. Viable CD45.1-B220+CD138-GL7-IgD-S1pr2+ and CD45.1-B220+CD138-GL7-IgD-S1pr2-B cells were sorted. Viable CD45.1+ B220+CD138-GL7-IgD-B cells were sorted as positive controls for anti-ssDNA autoantibody production. Sorting was performed on the SH-800Z cell sorter (Sony Biotechnology). 5000 B cells were then cultured in the presence of 50,000 sorted B220-cells and with 1ug/mL R848 (tlrl-r848, InVivoGen) and anti-CD40 (clone FGK4.5, BE0016-2, BioXCell) for 6 days. Anti-ssDNA autoantibody production was then assessed by ELISA.

### Immunization

To generate (transient) foreign-Ag-elicited GCs, mice were immunized intraperitoneally with 100 µg of chicken gamma globulin (Rockland Immunochemicals) precipitated in an equal volume of Imject Alum (ThermoScientific). To generate chronic GCs, mice were immunized with sheep red blood cells (SRBC) twice. For SRBC immunizations, 10–15 mL of Sheep blood in Alsevers (Colorado Serum Company) was washed two times in HBSS before counting SRBC using a Neubauer hemacytometer. Mice received a single injection of 2E8 SRBC in HBSS i.p followed by a boost of 2E8 SRBC 7 days later.

### Confetti analysis

Analysis was performed as described in the manuscript of S.Degn et al. (*7*). Briefly, acquired stacks were rendered in Fiji (ImageJ). Counting was performed on representative z-plane from stacks acquired at 5-μm steps through the GC, with multiple GCs per mouse. Raw counts were converted to relative frequencies for each observable color to simplify analysis. Clonal dominance was calculated as the frequency of the most dominant clone/color at a given time point. The clonal divergence score is calculated at a given time point by comparing the observed distribution of colors at time t to the expected frequencies resulting from random recombination, as reported previously (*9*). Expected frequencies for each of 10 possible colors were derived using explant data from day 3 post-tamoxifen treatment, where color distribution most closely mimics random recombination and before any selection events. Descriptors of GC behavior and clonality were graphed using GraphPad Prism 8.

### Flow cytometry

Spleens and LNs were harvested into ice-cold FACS buffer and mechanically dissociated using pestles in 1.5 ml Eppendorf tubes. Samples were filtered through 70-µm cell strainers (Corning) and spun down at ∼200 g for 5 min. LN samples were resuspended in FACS buffer. Spleen samples were resuspended in RBC lysis buffer (155 mM NH4Cl, 12 mM NaHCO3, 0.1 mM EDTA), incubated for 2–3 minutes then spun down as before and resuspended in FACS buffer. Samples were added to wells of 96-well round-bottom plates, spun down, and resuspended in 100 µl staining mix (appropriate Ab and viability dye cocktail in FACS buffer). Staining was performed for 30 min on ice in the dark, followed by the addition of 150 µl of wash buffer. Plates were subsequently centrifuged at 200 g for 5 min, and supernatants were flicked out of the plates. For two-step staining procedures, secondary staining mix was added (appropriate secondary Ab mix and viability dye in FACS buffer) and the process was repeated. When appropriate, cells were permeabilized for 30 min on ice and subsequently washed with perm/wash buffer. Appropriate intracellular Ab staining mix in perm/wash buffer was added for 30-45min on ice and subsequently washed. Following the last wash, samples were resuspended in 200 µl FACS buffer and transferred to FACS tubes. Flow cytometric analyses were performed on a FASCSCanto2, with 8-color and 10-parameter analytical capabilities.

### Immunofluorescence confocal microscopy

Freshly harvested tissue was put in 4% PFA for several hours on ice and subsequently embedded in OCT (TissueTek) and immediately frozen at −80°C. Tissue blocks were equilibrated at cutting temperature (−16 to −20°C depending on tissue type), and 10-15 µm thick sections were cut on a cryostat. Tissue sections were mounted on SuperFrost+ slides (Fisher Scientific) and fixed using ice-cold acetone. Slides were then rinsed with PBS and incubated with block/perm buffer (PBS, 2% FBS, 0.1% NaN3, and 0.1% Triton-X100) for 30 min. This was followed by incubation with primary Ab mixture in staining buffer (PBS, 2% FBS, 0.1% NaN3), overnight at 4°C. For two-step staining procedures, the slides were washed 3 times with PBS, 0.01% Tween-20. Then secondary Ab mixture in staining buffer was added and incubated for 2 hrs at RT. At the end of either one-or two-step staining procedures, slides were washed once with staining buffer for 5 min, then 3 times for 5 min with PBS, 0.01% Tween-20. Slides were spot-dried, then mounted in Fluoro-Gel (Electron Microscopy Sciences), and coverslipped. Imaging was performed using a Fluoview FV1000 inverted Olympus IX 81 confocal microscope, equipped with 6 laser lines (405, 457, 488, 515, 559, 635 nms) and 4 fluorescence + 1 transmission detectors (PMTs).

For each mouse, 2 non-consecutive sections of the spleen (separated by at least 50mM during cryosectioning) were stained and imaged on the FV3000R resonant scanning confocal microscope (Olympus). Images were acquired at 30x magnification with at least 4 individual fields of view per section to cover the entire area. To visualize entire spleens for downstream analysis, manual tiling of individual fields of view was carried out for each section. To determine the frequency of GCs with 564Igi and WT, 564Igi only or WT only ‘residents’, each section was manually examined and assigned to one of the groups. In parallel, GCs and white pulp areas were enumerated. GC area was determined using the ‘measure’ function in Fiji. For each mouse, the plotted values represent the average of the two non-consecutive splenic sections.

### Additional procedures

Recombination of the Rosa26Confetti allele in AicdaCreERT2 mice was induced by a single gavage of 10 or 15 mg of tamoxifen (Sigma) dissolved in sunflower oil at 20 or 30 mg/ml. For the transfer of S1PR2CreERT2 TdTomato B cells into 564Igi AicdaCreERT2 eYFP mice, tamoxifen gavage was performed on day 3 and day 5 post-adoptive transfer; mice were sac’ed on day 7 post-adoptive transfer. For the transfer of S1PR2CreERT2 TdTomato B cells into 564IgiHomo (CD45.1) mice for anti-ssDNA IgG analysis, a tamoxifen-containing diet was started 21 days prior and continued during adoptive transfer.

### Multiphoton imaging

Spleens were harvested at different times post-tamoxifen. Thick transversal sections (∼1 mm) were manually cut from spleens using surgical scissors. Spleen sections were placed in PBS in vacuum-grease chambers on microscope slides. Chambers were coverslipped and kept on ice before imaging. All imaging was performed on an upright Olympus FV1200 MPE multiphoton system microscope fitted with either a 20× 0.95NA Plan water-immersion objective or a 25× 1.05NA Plan IR optimized water-immersion objective, a MaiTai HP DeepSee Ti-Sapphire laser (Spectraphysics), and 4 non-descanned detectors (2 GaAsP and 2 regular PMTs). Imaging of Confetti alleles was performed using λ=940 nm excitation. Fluorescence emission was collected in three channels, using the following filter sets: a pair of CFP (480/40 nm) and YFP (525/50 nm) filters, separated by a 505 nm dichroic mirror, for CFP/GFP/YFP detection, and a third filter (605/70 nm) for RFP detection.

### Autoantibody array

MILLIPLEX MAP Human Autoimmune Autoantibody Panel (HAIAB-10K, Millipore Sigma) was used following manufacturer’s instructions. Due to supply chain issues, ribosomal P antigen was not included in the original kit, and the assay was performed against 19 autoantigens instead of the original 20. Serum was collected from whole blood by allowing clotting at room temperature for 30 minutes, followed by 2 centrifugation steps at 4°C: 1000g for 10 minutes, and 2700g for 2 minutes, collecting the supernatant after each step. BPI (Surmonics, Cat. A192), Ro60 (Surmonics, Cat.A174), and APRIL (Peprotech, Cat. 310-10C) were added to the assay. For anti-IgG2c (Cat 1079-09, Southern Biotech) and IgG-PE (Cat 1030-09, Southern Biotech), the amount of protein used for coating the beads was 0.005 nmoles; for the other proteins BPI, APRIL, Ro60 the concentration was 0.5 nmoles. For all beads, the calculations were made to have at least 400 beads of each protein (region IDs) at the start of each assay. Serum from 3 parabiont pairs of each type was included in the autoantigen array.

### Autoantibody measurement

*Anti-single-stranded DNA autoantibody measurement by ELISA.* ssDNA salmon DNA 20ul/ml (D9156, Sigma) was diluted in 5% DMSO in carbonate/bicarbonate coating buffer and incubated at 95C for 10 minutes. It was then coated on a 96-well ELISA plate and incubated overnight. The plate was washed x3 with wash buffer (0.01% Tween20 in PBS), then blocking buffer (5%BSA, 1%FCS in PBS) was added at 37C for 1h. The plate was washed x3 with wash buffer and sera (diluted 1:20 in 10% blocking buffer in PBS (dilution buffer)), added to the plate, and incubated at 4C overnight. Cell culture supernatants were added undiluted. The plate was washed x3, goat anti-mouse IgG2c-HRP (Southern Biotech, 1079-05) was diluted in dilution buffer and the plate was incubated at 37C for 1h. After washing the plate x3, TMB substrate was added for 15-20 minutes and the reaction was stopped with H2SO4. The 450nm absorbance was measured on the X Mark microplate spectrophotometer (Biorad). C11 (564Igi antibody derived from a hybridoma) was used to standardize absolute levels of anti-ssDNA antibody.

### Hep-2 Immunofluorescence assay

ANA (Hep-2) slides (AN-1012, MBL International) and anti-neutrophil cytoplasmic antibody (ANCA) slides (29419, Biorad) were removed from pouches and allowed to equilibrate at room temperature. Sera were diluted (1:20) in PBS, and 30uL was added to each well for 1h at room temperature. Slides were washed twice with blocking buffer (0.1% Tween20, 0.5% BSA in PBS). Secondary antibody goat anti-mouse IgG2c-AF488 (1079-30, Southern Biotech) and (CTB)-AlexaFluor555 (C34776, Thermo Fisher Scientific) were diluted in blocking buffer, added to the slides, and incubated for 1h at room temperature. Slides were washed twice with blocking buffer and mounted with DAPI Fluoromount-G mounting medium (0100-20, Southern Biotech).

### Image analysis

Images were quantified using CellProfiler. For HEp-2 slides, quantifications were done based on the DAPI (nucleus) and Phalloidin staining. Cell outlines were detected using the phalloidin staining and as a function of the detected nuclei. A cytoplasmic mask was generated by subtracting the nucleus mask from the cell mask. Each bar represents a mean of individual objects (for instance a mean of MFI from 20 nuclei).

### Statistical analysis

Statistical tests are described in the figure legends. Data were analyzed using GraphPad Prism software (GraphPad 8, San Diego, CA). No statistical methods were used to predetermine the sample size before the experiments.

## Supplementary Materials

Figs. S1 to S4

## Acknowledgments

We thank E.M. Carroll for administrative and technical support and all the members of the M.C.C. lab for help with experiments and feedback on the manuscript. Imaging was performed at the Program in Cellular and Molecular Medicine (PCMM) microscopy core, with technical support from Harry Leung. Flow cytometry and cell sorting were performed at the PCMM flow cytometry core, with technical support from Jodene K. Moore.

## Author contributions

Conceptualization: MCC, TB Methodology: MCC, TB, KO, CC, CEP Investigation: TB, KO, SR Formal analysis: TB, KO Visualization: TB Funding acquisition: MCC TB Project administration: MCC Supervision: MCC Writing – original draft: TB Writing – review & editing: MCC, TB, KO, CC

## Data and materials availability

All data are available in the main text or the supplementary materials.

---

**Supplemental figure 1.**
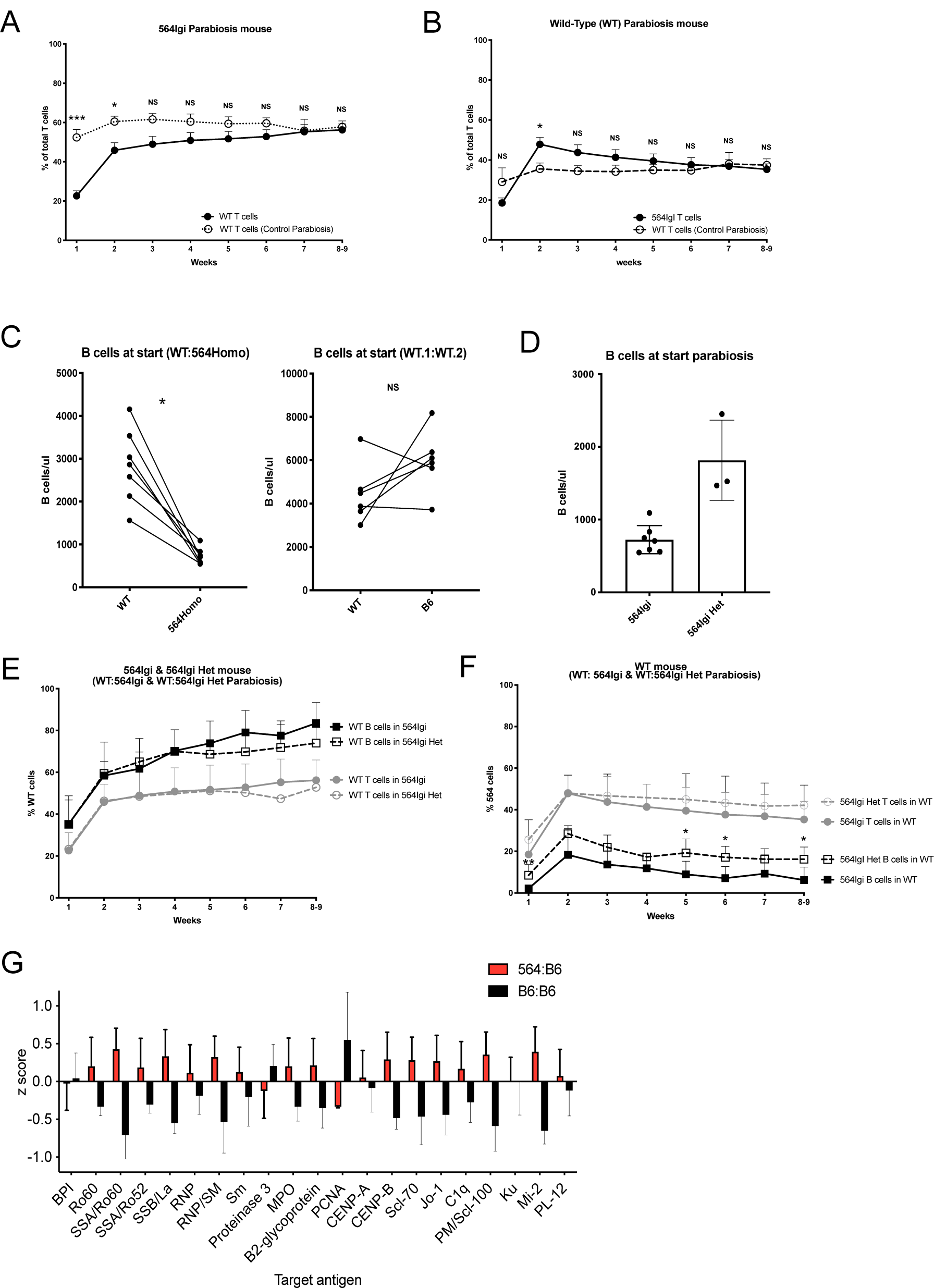
Exchange of T cells and presence of autoantibodies in parabiosis. A. Proportion of wild-type (WT) T cells in the circulation of 564Igi parabiosis partner (closed circles) or WT parabiosis partner (control parabiosis, open circles) over time. B. Proportion of 564Igi (closed circles) or WT (control parabiosis, open circles) T cells in the circulation of WT parabiosis partner mouse over time. C. Absolute number (cells/ul) of B cells at the start of parabiosis surgery in circulation for WT with 564Igi parabiosis (left graph) and WT with WT control parabiosis (right graph). D. Absolute number (cells/ul) of B cells in the circulation of 564Igi and 564Igi heterozygous mice. E. Proportion of wild-type (WT) cells in the circulation of 564Igi or 564Igi heterozygous (564Igi het) parabiosis partner for B cells (squares) and T cells (circles), over time (n=5 pairs of 564heterozygous with WT). F. Proportion of 564Igi or 564Igi heterozygous (564Igi het) cells in the circulation of WT parabiosis partner for B cells (squares) and T cells (circles), over time (n=5 pairs of 564heterozygous with WT). G. IgG2c (WT) serum auto-reactivity in 564Igi parabiosis and control parabiosis, z score is the difference of the average per specific reactivity. Significance is indicated as ***P<0.001, **P<0.01, *P<0.05 and P>0.05 (not significant (NS)). Number of measurements in S1A,1B as shown in figure 1B,1C. Number of measurements in S1E. WT B cells in 564Het mouse, week 1,2,3,5,8-9 n=5, week 4 n=1, week 6 n=4, week 7 n=3. WT T cells in 564Het same as for WT B cells in 564Het but at week 4 n=0. Number of measurements in S1F. 564Het T cells in WT mouse, week 1,2,3,5,6,7,8-9 n=5, week 4 n=0. For 564Het B cells in WT mouse, week 1,2,3,5,6,7,8-9 n=5, week 4 n=1. S1G, serum of 6 parabiosis pairs. By Mann-Whitney test fig S1A, S1B, S1H, S1I. Wilcoxon paired test fig S1C, S1E, S1F.

**Supplemental figure 2.**
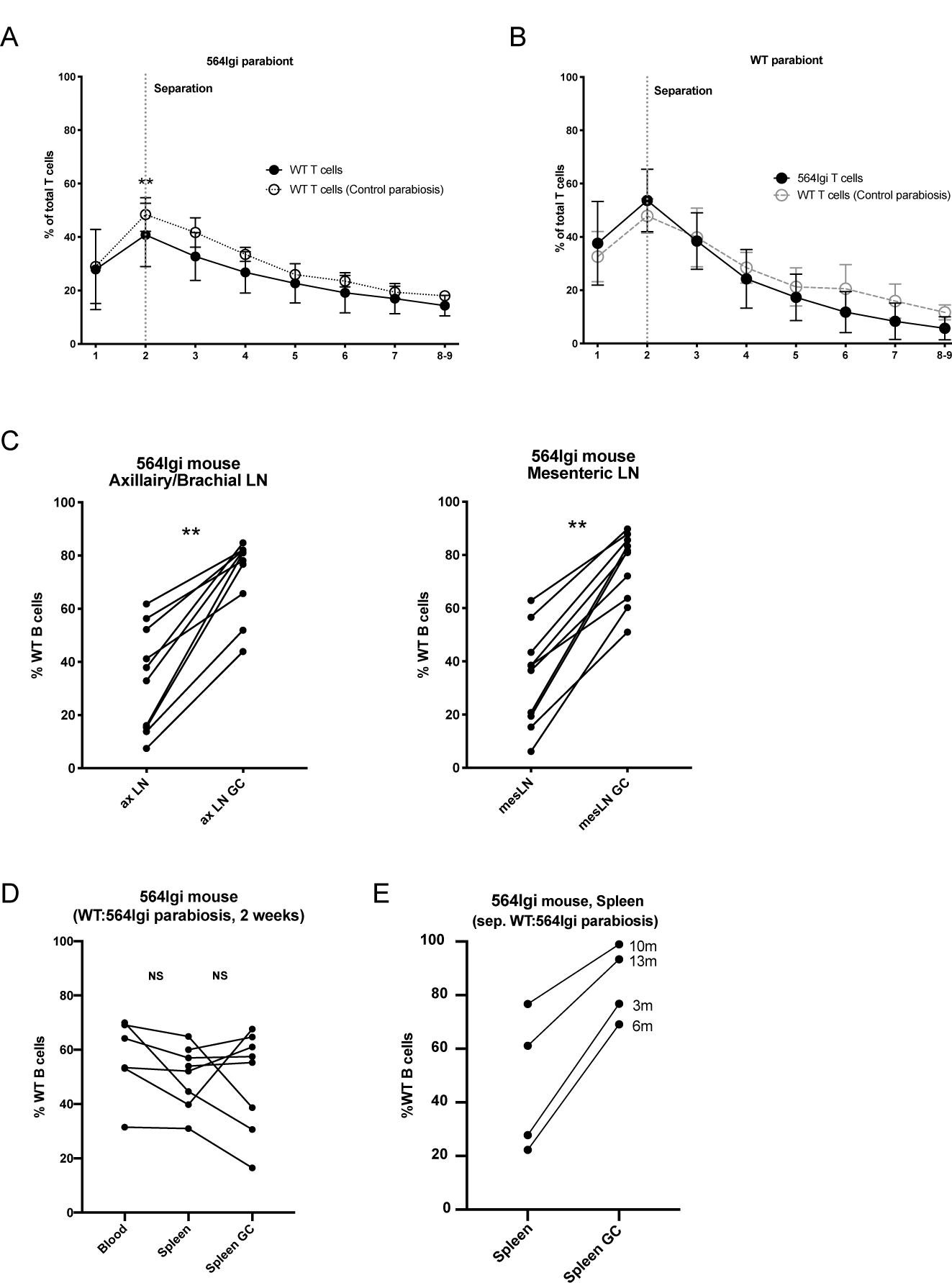
WT B cells persist in the GC of lymph nodes and spleen. A. Proportion of wild-type (WT) T cells in the circulation of 564Igi (black circles) or WT (control parabiosis, open circle) parabiosis partner mouse over time. B. Proportion of 564Igi (closed squares) or WT (Control parabiosis, open squares) T cells in the circulation of WT parabiosis partner mouse over time C. Proportion of WT B cells in axillary/brachial (first graph, n=10) and mesenteric (second graph, n=10) lymph node (LN) of 564Igi parabiosis partner. D. Proportion of WT B cells in circulation, splenic B cell (spleen) and splenic GC B cell (Spleen GC) compartment in 564Igi parabiosis partner, 2 weeks post-parabiosis surgery (n=8). E. The proportion of WT B cells in 564Igi parabiosis partner in total splenic B cell population or within the splenic germinal center B cell population, after 2 weeks of parabiosis and described time of separation (n=4). Figure S3A, S3B: same number of mice as figure 3B,3C respectively. Significance is indicated as **P<0.01, and P>0.05 (not significant (NS)). By Mann-Whitney test fig S3A, S3B. Wilcoxon paired test fig S3C, S3D, S3E (Blood vs tissue, tissue vs tissue GC)

**Supplemental figure 3.**
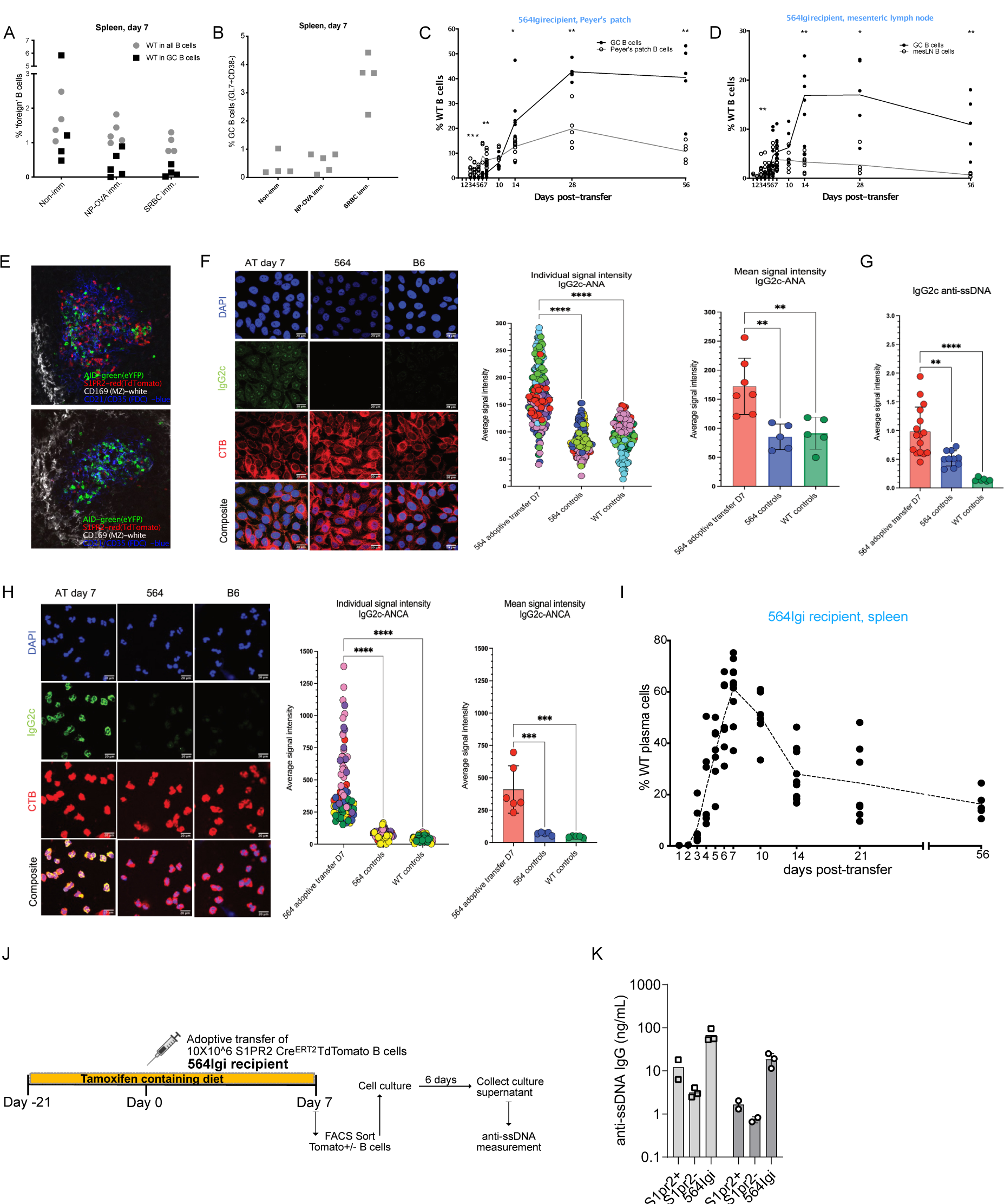
WT B cells enter the autoreactive center and generate auto-antibodies. (A) The proportion of transferred WT B cells within the spleen (gray circle) and splenic germinal center (black square) in non-immunized wild-type B6 mice (left), NP-OVA immunized WT B6 mice (middle), sheep red blood cell (SRBC)-immunized WT B6 mice (right) (B) The proportion of germinal center B cells (B220+ GL7+ CD38-) in WT mice (left), NP-OVA immunized mice (middle), SRBC-immunized mice (right) (C) Proportion of WT B cells within B cell pool of Peyer’s patches (grey line, open circle) and within germinal center B cell pool (black line, closed circle) over time. (D) Proportion of WT B cells within B cell pool of mesenteric lymph node (grey line, open circle) and within germinal center B cell pool (black line, closed circle) over time. (E) Transfer of S1PR2-Cre-ert2 TdTomato B cells into 564Igi AID-Cre-ert2 YFP mouse, 7 days post-transfer, S1PR2+ WT B cells (Red), AID-YFP+ 564Igi B cells (Green), FDC (CD21/23, Blue), marginal zone (CD169, Grey) (F) ANA detection from sera of 564Igi mice 7 days after WT B cells transfer. Representative images of ANA Hep-2 slides are shown with quantifications of signal intensity to cytoplasmic, nuclear and/or nucleolar antigens. The scatter-dot plot showed individual points from each individual data points collected (left) and the bar chart visualized the mean signal intensity in each group of mice. The scale bars represent 20mm. (adoptive transfer n=7, 564Igi control n=5, WT control n=5) (G) ANA detection from sera of 564 Igi mice 7 days after WT B cells transfer. Images of ANCA slides are shown with quantifications of signal intensity to cytoplasmic neutrophilic antigens. The scatter-dot plot showed individual points from each individual data points collected (left) and the bar chart visualized the mean signal intensity in each group of mice. The scale bars represent 20mm. ((adoptive transfer n=6, 564Igi control n=5, WT control n=5) (H) Analysis of serum titers of IgG2c anti-ssDNA. (Adoptive transfer n=16, 564Igi control n=10, WT control n=6) (I) Proportion of WT plasma cells within splenic plasma cell pool (B220+CD138+) of 564Igi mice over time after WT B cell adoptive transfer (J) Schematic experimental setup to analyze antibody reactivity of GC-experienced WT B cells Anti-ssDNA IgG from sorted GC-experienced (S1PR2+) and GC-inexperienced (S1PR2-) WT B cells, and 564Igi B cells (positive control), n= 2 mice, 2-3 replicates. These data represent at least two independent experiments. Statistical significance was measured by one-way ANOVA (**p < 0.01, ***p < 0.001, ****p < 0.0001). Fig3C,D with Mann-Whitney test at each individual timepoint.

**Supplemental figure 4.**
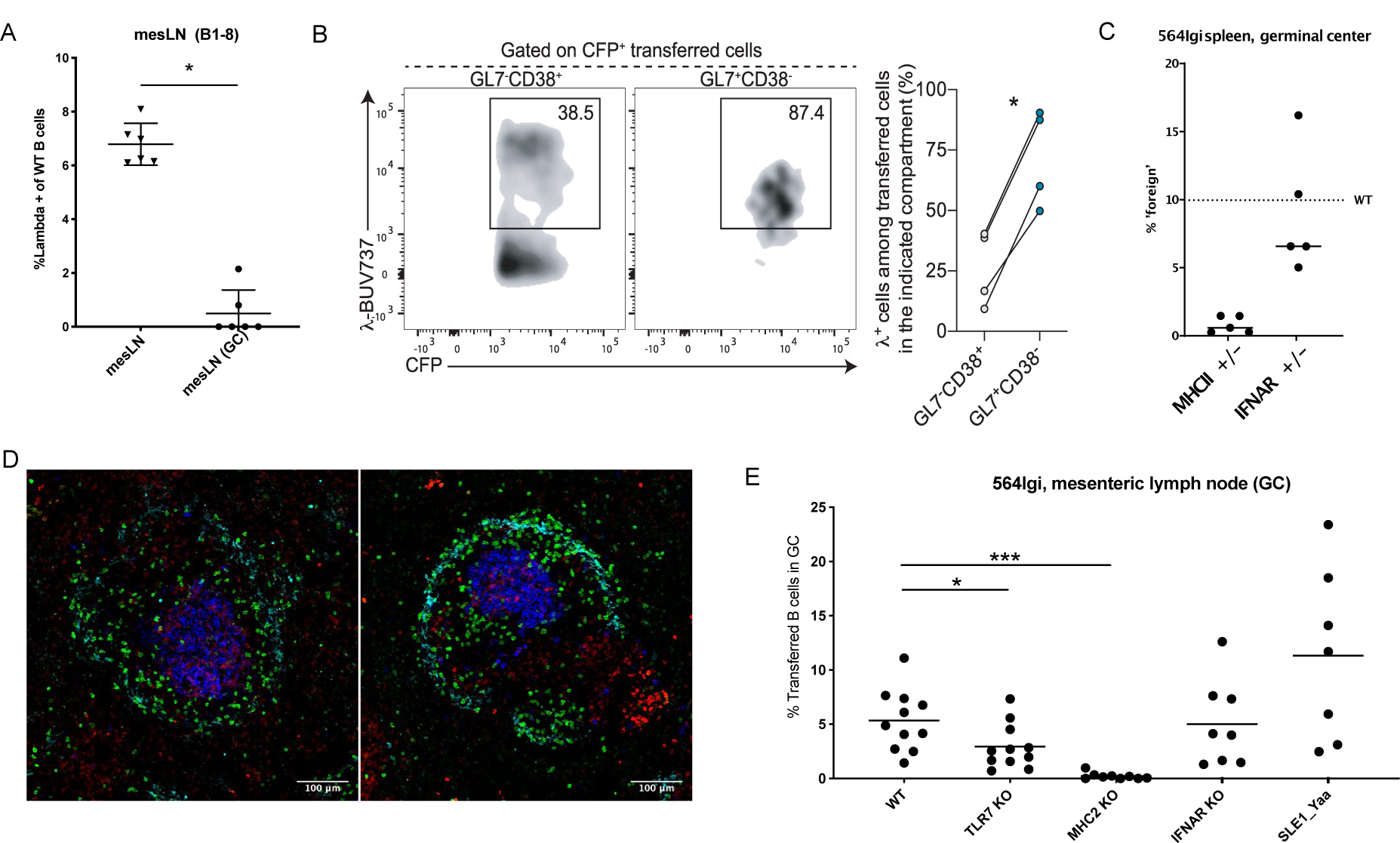
Molecular requirements for wild-type B cell entry and participation in germinal centres of mesenteric lymph node. A. Proportion of Lambda-positive (NP-specific) B cells within WT B cell transferred population within the mesenteric lymph node (mesLN), mesLN germinal center (mesLN(GC)) compartment, day 7 post-adoptive transfer. B. Representative flow cytometry plots (left) and summary scatter plot (right) with the frequency of NP-specific (lambda+) B cells among GL7-CD38+ and GL7+CD38-transferred cells (CFP+) in the spleens of recipient mice on day 7 post transfer C. Proportion of haploinsufficient MHCII and IFNAR B cells within splenic germinal center of the 564Igi mouse, day 7 post-adoptive transfer. Dashed line highlights mean value of WT adoptive transfer for germinal center (GC), as in Fig 5C. D. Representative confocal image of adoptive transfer of MHC2-KO B cells into 564Igi mouse, 7 days post-transfer, MHC2-KO B cells (CD45.2, Green), Plasma cells (CD138, Red), Germinal Center (GL7, Blue), marginal zone (CD169, Cyan). E. The proportion of the different transferred B cells within the germinal center of the mesenteric lymph node, day 7 post-adoptive transfer. Significance is indicated as ***P<0.001, **P<0.01, *P<0.05. Fig C IFNAR+/-transfer n=5, MHCII+/-n=5. Fig D: WT transfer n=11, MHC2 KO transfer n=9, TLR7 KO transfer n=11, SLE1-Yaa n=7, IFNAR KO transfer n=8. Statistics: Man-Whitney test S4A,B,C,E (compared to WT).

